# Biphasic effects of IL-27 during *Staphylococcus aureus* implant-associated osteomyelitis in mice

**DOI:** 10.1101/2021.05.20.444931

**Authors:** Yugo Morita, Anthony M. Franchini, John R. Owen, John C. Martinez, John L. Daiss, Karen L. de Mesy Bentley, Stephen L. Kates, Edward M. Schwarz, Gowrishankar Muthukrishnan

**Affiliations:** Center for Musculoskeletal Research, University of Rochester Medical Center, Rochester, NY, USA; Department of Environmental Medicine, University of Rochester School of Medicine and Dentistry, Rochester, NY, USA; Department of Orthopaedic Surgery, Virginia Commonwealth University, Richmond, VA, USA; Department of Orthopaedics, University of Rochester Medical Center, Rochester, NY, USA; Department of Pathology and Laboratory Medicine, University of Rochester Medical Center, Rochester, NY, USA

**Keywords:** interleukin-27, *S. aureus*, osteomyelitis, staphylococcal abscess, osteolysis, host-pathogen interactions

## Abstract

Interleukin-27 is a pleiotropic cytokine whose reported functions during bacterial infections are debated as an area of active research. To address this, we investigated the role of IL-27 signaling during *Staphylococcus aureus* osteomyelitis. Clinically, we observed elevated serum IL-27 levels (20-fold higher, p<0.05) in patients with *S. aureus* osteomyelitis compared to uninfected patients undergoing elective total joint replacement. Remarkably, IL-27 serum levels immediately following septic death were 60-fold higher vs. uninfected patients (p<0.05), suggesting that IL-27 may be a biomarker of end-stage infection and/or cytokine storm. To test this, we hypothesized that IL-27 mediates bacterial clearance during the acute phase of *S. aureus* osteomyelitis, and subsequently suppresses inflammation to prevent cytokine storm and osteolysis during chronic infection. In mice, we observed that systemic IL-27 delivery by a recombinant adeno-associated viral vector (rAAV-IL-27) ameliorates surgical site soft tissue infection and peri-implant bone loss during the establishment of implant-associated *S. aureus* osteomyelitis. This effect was not observed in IL-27 receptor α knock-out mice, suggesting a direct role of IL-27/IL-27R signaling on immune and bone cell functions. Examination of IL-27-mediated immune responses via transcriptome analyses of infected tibiae demonstrated that IL-27 is a biphasic cytokine with IL-27/IL-27R activating immunostimulatory responses including Th17, IL-2, TLR, and iNOS signaling early, and subsequently suppressing these pathways during chronic infection. Ex vivo confirmation using murine macrophages revealed that IL-27 co-stimulates TLR signaling to increase the production of nitric oxide, and immunomodulatory cytokines such as IL-10, IL-21, IL-31, and TNF-β, but is not a chemokine.

**Author Summary:** *Staphylococcus aureus* is the most common pathogen in orthopaedic infections, and hard-to-treat (MRSA) strains cause >50% of these infections. Thus, there is an urgent need to develop immunotherapies to treat these life-threatening *S. aureus* infections. Currently, the role of multifunctional IL-27 on *S. aureus* osteomyelitis is unknown. In a clinical study, we observed that IL-27 is an important biomarker for identifying *S. aureus* osteomyelitis patients, and that elevated serum IL-27 levels correlated with adverse clinical outcomes, such as septic death. In our efforts to uncover the underlying mechanisms, we reveal that IL-27 is a biphasic cytokine, activating proinflammatory immune pathways, including Th17 responses, early during acute *S. aureus* osteomyelitis, and subsequently repressing them during the chronic phase to prevent cytokine storm and bone damage. These results indicate that immune modulation of IL-27/IL-27R signaling could be a viable therapeutic strategy in mitigating *S. aureus* osteomyelitis.

## Introduction

Deep bone infections continue to be the bane of orthopaedic surgery, with infection rates essentially remaining at 1-2% for elective surgery over the past 50 years, despite significant medical advances [1–3]. *Staphylococcus aureus* is the major pathogen in orthopaedic infections. It is responsible for causing 10,000-20,000 prosthetic joint infections (PJI) annually in the United States alone [4, 5] and 30-42% of fracture-related infections (FRI) [6, 7]. Unfortunately, these difficult-to-treat *S. aureus* bone infections are associated with poor clinical outcomes and high recurrence rates following revision surgery [8, 9]. With increasing methicillin-resistant *S. aureus* (MRSA) osteomyelitis incidence rates, and emerging strains with pan-resistance [10, 11], there is an urgent need for novel immunotherapies to supplement existing antibiotic therapies.

*S. aureus* causes the most lethal form of human sepsis with a 10% mortality rate, and a catastrophic outcome of osteomyelitis is death due to sepsis and multiple organ failure [12, 13]. The mechanisms behind *S. aureus* osteomyelitis-induced sepsis are largely unknown. Interestingly, several studies have reported elevated serum IL-27 levels during sepsis, suggesting that IL-27 could potentially be a diagnostic biomarker of sepsis [14–19]. IL-27 is a heterodimeric cytokine belonging to the IL-12 cytokine family and is mainly produced by antigen presenting cells such as macrophages, monocytes, and dendritic cells [20, 21]. It is composed of IL-27p28 and EBI3 subunits, and signals through a heterodimeric cell surface receptor composed of IL-27 receptor α (IL-27Rα) and gp130 [22, 23]. Like IL-12, it signals mainly through the JAK-STAT intracellular pathway and plays a central role in multiple immune regulation activities. It downregulates Th17 differentiation, stimulates regulatory T cell formation, and directly modifies CD4+ T cell effector functions to induce anti-inflammatory IL-10 [20, 21, 24, 25]. Studies involving cecal ligation and puncture (CLP)-induced bacterial sepsis and *S. aureus* pneumonia following influenza demonstrated that IL-27 regulates enhanced susceptibility to infection by attenuating Th17 immunity and promoting IL-10 induction [26, 27]. These studies highlight the importance of IL-27 in immune suppression. On the other hand, IL-27 has been reported to promote proliferation and differentiation of hematopoietic stem cells [28], increase production of proinflammatory cytokines by monocytes [29, 30], and induce Th1 differentiation [31]. Currently, the role of IL-27 in host immunity during *S. aureus* osteomyelitis is unknown. Here, we tested the hypothesis that IL-27 is a biphasic cytokine that enhances bacteria killing via promoting inflammation early during acute *S. aureus* osteomyelitis, and subsequently suppresses inflammation during chronic bone infection to prevent cytokine storm and osteolysis. Consistent with this theory, we report that IL-27 is induced in patients with *S. aureus* osteomyelitis, and elevated serum IL-27 correlated with septic death in these patients. Examining IL-27’s role in mice revealed that this cytokine is crucial for carefully balancing the host immunostimulatory and immunosuppressive responses during *S. aureus* osteomyelitis.

## Results

### *S. aureus* infection induces IL-27 secretion in patients and in mice

To better understand host immune responses against *S. aureus* osteomyelitis, we analyzed sera from healthy people, orthopaedic patients with culture-confirmed *S. aureus* bone infections, and patients who died from septic *S. aureus* osteomyelitis. Serum IL-27 levels were significantly elevated in infected patients compared to uninfected individuals (20-fold higher, p<0.05). Remarkably, IL-27 levels immediately following septic death were 60-fold higher (**Fig. 1A**, p<0.05), suggesting that IL-27 could be an essential biomarker for *S. aureus* osteomyelitis-induced septic death. Indeed, formal analyses of IL-27 as a diagnostic biomarker using receiver operator characteristic (ROC) curve analysis revealed a high area under the curve (AUC) of 0.922 (**Fig. 1B**, p<0.0001). We also evaluated whether *S. aureus* infection directly induces IL-27 production in murine macrophages in vitro. Interestingly, in both RAW 264.7 macrophages and murine bone marrow-derived macrophages, *S. aureus* induced significant IL-27 secretion 24 hours post infection in M0, M1, and M2 murine macrophages (**Fig. 1C-D**, p<0.05). Collectively, these data indicate an important role for IL-27 in host immunity against *S. aureus* infections.

**Figure 1.**
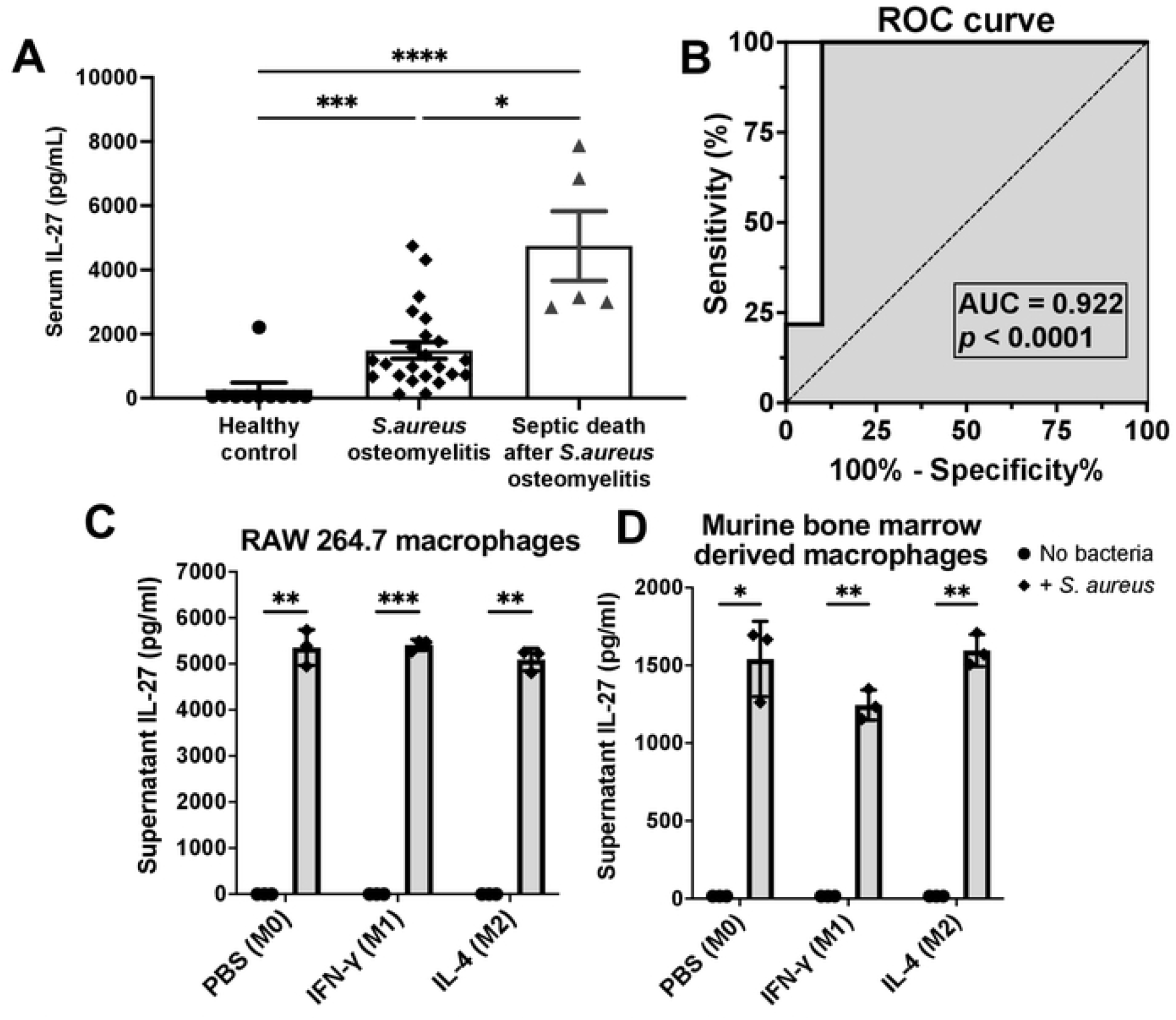
IL-27 levels are elevated in patients and in mice with *S. aureus* infection. **A)** Serum samples were collected from healthy people (n=10), orthopaedic patients with culture confirmed *S. aureus* bone infections (n=23), and patients who died from septic *S. aureus* osteomyelitis (n=5). Serum IL-27 levels were determined via Luminex assay, and the data for each sample is presented with the mean +/− SEM for the group. **B)** The Luminex data were utilized to perform a receiver operating characteristic (ROC) curve, and the area under the curve (AUC) between controls and infected patients is presented. Note that serum IL-27 levels are highly diagnostic of *S. aureus* osteomyelitis. The dashed line represents a non-discriminatory test with equal sensitivity and specificity. In vitro cultures of (**C**) RAW264.7 cells and (**D**) primary murine bone marrow-derived macrophages were differentiated with PBS, IFN-γ (50ng/ml) or IL-4 (20ng/ml) to generate M0, M1 and M2 cells respectively, and then exposed to *S. aureus* USA300 (MOI = 10). These cultures were incubated for 24 hours, and IL-27 levels in the supernatants were assessed via ELISA. The data from each experiment are presented with the mean +/− SD for the group (n=3) (\**p*<0.05, \*\**p*<0.01, \*\*\**p*<0.001 \*\*\*\**p*<0.0001, ANOVA).

### Systemic IL-27 delivery inhibits draining abscess formation and bone osteolysis during establishment of *S. aureus* osteomyelitis

Having established an association between IL-27 and *S. aureus* osteomyelitis in patients and mice, we next examined if IL-27 mediates bacterial clearance during *S. aureus* osteomyelitis using our well-established murine model of osteomyelitis [32–36]. Mice were challenged with bioluminescent MRSA (USA300 LAC::lux) via transtibial implantation of a contaminated stainless-steel implant following intramuscular injection of rAAV-IL-27 or adeno-associated virus expressing recombinant GFP (rAAV-GFP, control). Before the in vivo infection experiment, we confirmed exogenous IL-27 expression in mouse sera out to day 24 following intramuscular injection of rAAV-IL-27 (**Fig. 2A**). While rAAV-IL-27 treatment did not show an effect on in vivo *S. aureus* growth as assessed by bioluminescent intensity (BLI) (**Fig. 2B**), rAAV-IL-27 treated mice showed greater body weight recovery following septic-surgery compared to rAAV-GFP treated animals (**Fig. 2C**). Remarkably, rAAV-IL-27 treated animals showed much smaller draining abscess formation at the site of bone infection (**Fig. 2D**). Ex vivo CFU analyses confirmed that the bacterial load in surgical site soft tissues was significantly lower in rAAV-IL-27 treated mice (**Fig. 2E**). Moreover, high-resolution μCT demonstrated that peri-implant osteolysis was decreased in mice treated with rAAV-IL-27 compared to rAAV-GFP treated animals (**Fig. 2F**). These results demonstrate that IL-27 affects abscess formation and bone osteolysis. Interestingly, CFU quantification on the implants revealed similar bacterial loads between groups suggesting that systemic IL-27 treatment does not affect biofilm formation on the implant. Indeed, scanning electron microscopy (SEM) interrogation confirmed these findings (**Supplemental Fig. 1**).

**Figure 2.**
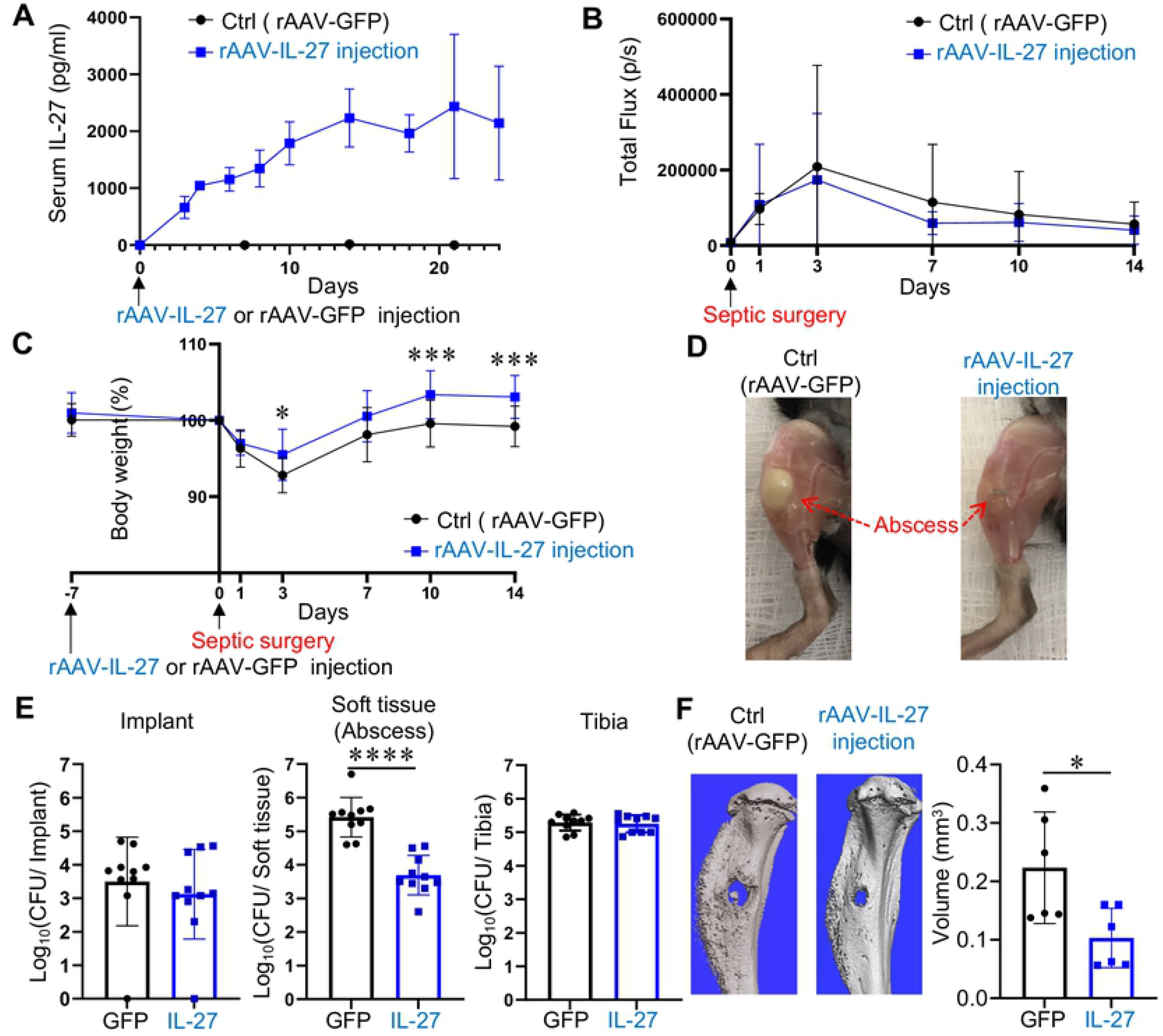
Systemic IL-27 ameliorates surgical site soft tissue infection and osteolysis during *S. aureus* implant-associated osteomyelitis. **(A)** 8-week-old female C57BL/6 mice received 0.5 × 10^12^ genome copies/mouse of rAAV-IL-27 (n=3) or rAAV-GFP (control, n=5) via intramuscular injection, and serum samples were collected longitudinally to assess IL-27 levels via ELISA. Exogenous IL-27 levels in sera are presented as the mean +/− SD. **(B-F)** A separate cohort of these mice were intramuscularly injected with rAAV-IL-27 or rAAV-GFP (n=16), and then challenged 7 days later with 5×10^5^ CFU of USA300 LAC::lux on a contaminated transtibial pin as described in Materials and Methods. Longitudinal BLI **(B)** and animal weight **(C)** were obtained on days 0, 1, 3, 7, 10 & 14, and the data are presented as the mean +/− SD (\**p*<0.05 on Day 3, \*\*\**p*<0.001 on Day 10 and 14, two-way ANOVA). **(D)** Photographs of the infected tibiae were obtained on day 14, and representative images of the large vs. small draining abscesses observed in the rAAV-GFP and rAAV-IL-27 treated mice respectively are shown. The mice were euthanized on day 14, and the infected tibiae were harvested for CFU and micro-CT analyses. **(E)** CFUs from the implant, soft tissue and tibia were determined, and the data for each tibia are presented with the mean +/− SD for the group (n=10, \*\*\*\**p*<0.0001, t test). **(F)** Representative 3D renderings of the extensive peri-implant osteolysis and reactive bone formation in rAAV-GFP vs. rAAV-IL-27 treated tibiae are shown with the volumetric bone loss in the infected tibiae. Data are presented for each tibia with the mean +/− SD for the group (n=6, \**p*<0.05, t test).

### Systemic IL-27 effects on *S. aureus* implant associated osteomyelitis in IL-27Rα^−/−^ mice

Two possible scenarios can lead to the observed suppression of *S. aureus* SACs and reduced bone osteolysis at the surgical site. IL-27 could be a chemokine attracting myeloid cells to the site of *S. aureus* infection. Alternatively, IL-27/IL-27R signaling pathway could extrinsically be inducing chemotaxis of innate immune cells to the infection site. First, we examined if IL-27 is chemotactic of myeloid cells. In vitro chemotaxis assay using granulocytic HL-60 cells revealed that IL-27 did not promote migration of granulocytes through the Boyden chambers (**Supplemental Fig. 2**). IL-27 was also not chemotactic of primary bone marrow-derived macrophages (data not shown). Next, to test whether IL-27/IL-27R signaling was inducing chemotaxis of immune cells to cause the observed phenotype, we repeated the in vivo *S. aureus* osteomyelitis experiments using IL-27 receptor α knock out (IL-27Rα^−/−^) mice. At 14 days post infection, body weight changes (**Fig. 3A**) and BLI (**Fig. 3B**) were similar between IL-27Rα^−/−^ mice treated with rAAV-IL-27 or rAAV-GFP. Most interestingly, ex vivo CFU on the implants, surgical site soft tissues, and tibia were similar in IL-27Rα^−/−^ mice (**Fig. 3D**). Furthermore, no difference was detected in draining abscess formation on these implants (**Fig. 3C**) and gross assessment of peri-implant osteolysis (data not shown) between groups. These data indicate that the effects of rAAV-IL-27 on *S. aureus* osteomyelitis in WT mice were due to IL-27/IL-27R signaling.

**Figure 3.**
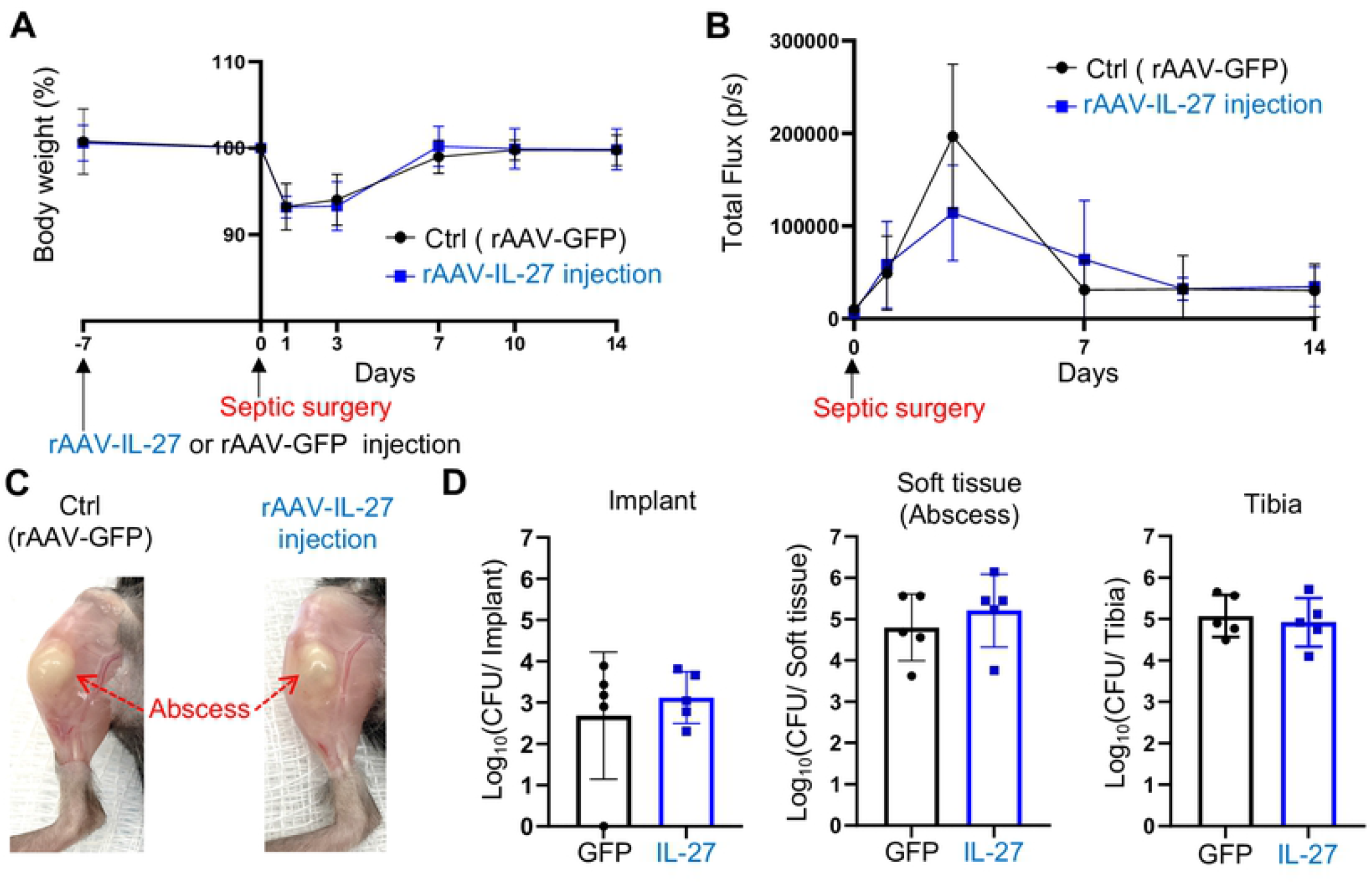
Absence of systemic IL-27 effects on implant associated osteomyelitis in IL-27Rα^−/−^ mice. Female IL-27Rα^−/−^ mice (C57BL/6 background) were intramuscularly injected with rAAV-IL-27 or rAAV-GFP and then challenged with a MRSA (USA300 LAC::lux) contaminated transtibial implant as described in Figure 2. Animal weight **(A)** and BLI **(B)** were obtained on days 0, 1, 3, 7, 10 & 14, and the data are presented as the mean +/− SD for the group (n=5). **(C)** Representative photographs obtained on day 14 post-surgery, illustrate similar large draining abscesses in both groups. **(D)** CFUs from the implant, surgical site soft tissue, and tibia were determined after euthanasia on day 14 post-op, and data from each tibia are presented with the mean +/− SD for the group (n=5). No differences were observed between the experimental groups.

### Identification of systemic IL-27 affected pathways during the establishment of implant-associated osteomyelitis

Next, we sought to elucidate the mechanism of IL-27/ IL-27R signaling effects on *S. aureus* osteomyelitis via unbiased gene expression studies. MRSA-infected mouse tibiae from rAAV-IL-27- and rAAV-GFP-treated groups were harvested on days 1, 3, 7, and 14 post-septic surgery and subjected to bulk RNA sequencing. The number of differentially (up-regulated or down-regulated) expressed genes (DEGs) on each day are shown in **Fig. 4B**. Venn diagram analyses of DEGs revealed *IL-27*, prostaglandin E synthase (*PTGES*), and sodium/myo-inositol cotransporter (*SLC5A3*) to be the common overlapping nodal points across all time points (**Fig. 4C**). Expectedly, IL-27 expression in the infected tibia was significantly up-regulated in mice treated with rAAV-IL-27 compared to rAAV-GFP at all time points (**Fig. 4D**), suggesting a positive feedback effect [37].

**Figure 4.**
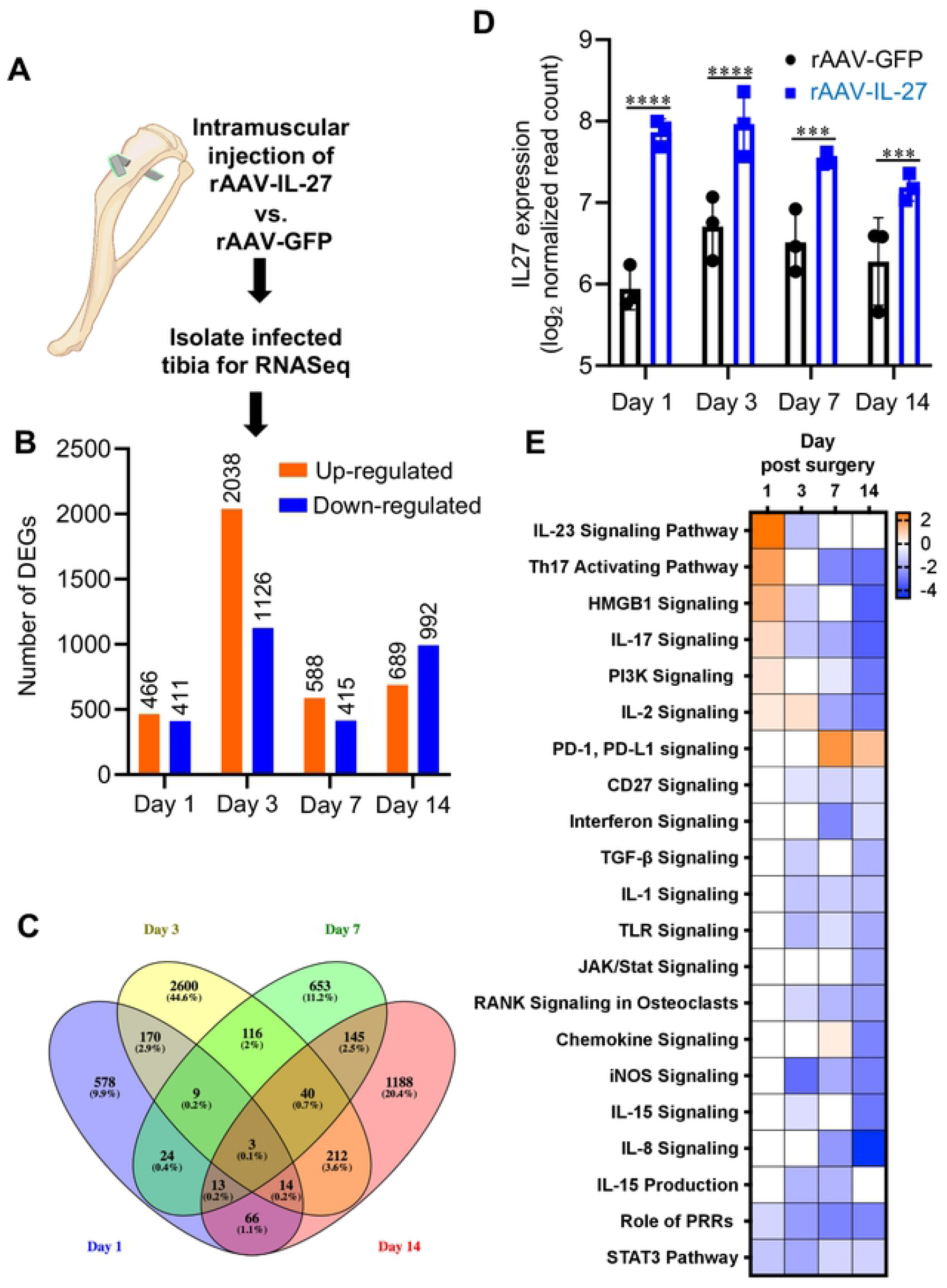
Identification of systemic IL-27 affected pathways during the establishment of implant-associated osteomyelitis via bulk RNA sequencing of *S. aureus-infected* tibia. **(A)** MRSA-infected tibiae were collected on days 1, 3, 7 and 14 post-septic surgery for bulk RNA sequencing (n=3), and differential gene expression between rAAV-IL-27 and rAAV-GFP treated mice were evaluated using DESeq2-1.22.1 R/Bioconductor package. **(B)** The number of significant differentially up-regulated and down-regulated genes in mice treated with rAAV-IL-27 on each day is shown. **(C)** Expression of *IL27* in infected tibiae was upregulated in mice treated with rAAV-IL-27, suggesting a positive feedback effect. (\*\*\*\**p*<0.0001 on days 1 & 3, \*\*\**p*<0.001 on days 7 & 14). **(D)** Venn diagram analyses showing the overlap of DEGs across days 1, 3, 7 and 14 post-septic surgery in in mice treated with rAAV-IL-27 vs. rAAV-GFP, respectively. **(E)** Ingenuity Pathway Analysis (IPA) was utilized to identify canonical pathways of DEGs between rAAV-IL-27 vs. rAAV-GFP treated mice over time. Significant association (\**p*< 0.05) were calculated based on the Fisher’s right tailed exact test. The orange and blue colored bars indicate predicted pathway activation or predicted pathway suppression in mice treated with rAAV-IL-27 vs. rAAV-GFP respectively (z-score). White bars indicate z-score at or very close to 0. Some pro-inflammatory/immune pathways including IL-17 signaling, Th17 activating pathway, IL-2 signaling were activated in mice treated with rAAV-IL-27 on day 1 following infection. On the other hand, these pathways and others (e.g.) were estimated to be suppressed at later time points. Moreover, immunosuppressive PD-1/PD-L1 signaling pathway was upregulated at later time points. These results indicate that IL-27 is a biphasic cytokine that activates pro-inflammatory/immune pathways early upon *S. aureus* infection, and then suppresses them late to prevent tissue damage and cytokine storm.

Additionally, Ingenuity Pathway Analysis (IPA) was performed to identify canonical pathways enriched between rAAV-IL-27 and rAAV-GFP treated animals (**Fig. 4E**). Examination of enriched pathways involved in innate and adaptive immunity revealed that pro-inflammatory immune pathways including IL-23 signaling pathway, Th17 activating pathway, IL-17 signaling, and IL-2 signaling were activated in rAAV-IL-27 treated mice during the acute phase of *S. aureus* osteomyelitis (day 1 post-surgery) compared to rAAV-GFP treated animals. Interestingly, these pathways were suppressed at later time points, especially on day 14, which represents the chronic phase of the disease. These data strongly indicate that IL-27 could be a biphasic cytokine, which activates pro-inflammatory immune pathways early upon *S. aureus* infection and suppresses them late to prevent tissue damage and cytokine storm.

### IL-27-mediated induction of pro-inflammatory cytokines early during *S. aureus* osteomyelitis and their down-regulation during chronic infection

RNAseq analyses revealed that genes associated with IL-23 signaling (**Fig. 5A**) and Th17 activation pathway (**Fig. 5B**) were significantly were up-regulated in mice treated with rAAV-IL-27 on day 1 post-surgery, compared to rAAV-GFP animals. These genes include *IL17A, IL-17F, IL-21,* and *IL12B.* We confirmed that IL-27-pretreated murine macrophages induce moderate production of pro-inflammatory cytokines such as IL-21, IL-31, and TNF-β early in response to *S. aureus* infection (**Table 1**). Of note, anti-inflammatory cytokine IL-10 was also modestly up-regulated in these macrophages suggesting a pleiotropic nature for IL-27. Remarkably, pro-inflammatory cytokine coding genes such as *IL17A*, *IL12A, TNF,* and *IL-6* were down-regulated later at day 14 during the chronic phase of infection (**Fig. 5A-B**). Utilizing the DEG data, we also assessed the top regulatory networks in IPA for these genes to provide further insight into the effects of differential gene expression in our dataset (**Supplemental Table 1**). Our analyses indicated that regulatory genes such as HSP90B1 and EGR2, up-regulated pro-inflammatory cytokine-coding genes *IL21, IL17A, IL12B, IL17F,* and *RORC* in rAAV-IL-27 treated mice. *RORC* encode the Th17 master transcription factor RORγt [38]. Additionally, transcriptome analyses revealed that immunostimulatory genes associated with Toll-like receptor (TLR) (**Fig. 5C**) and iNOS (**Fig. 5D**) signaling pathways were suppressed at later stages of *S. aureus* infection. However, their expression levels during the early infection phase (day 1) were equivocal. Indeed, we confirmed that combination of IL-27 and TLR agonist lipopolysaccharide (LPS) stimulation increased nitric oxide (NO^−^) production in primary macrophages, suggesting a co-immunostimulatory effect on TLR signaling (**Supplemental Fig. 3**). Collectively, these results indicate that IL-27 modulates immune homeostasis by promoting the production of pro-inflammatory cytokines early upon *S. aureus* infection and suppressing them late to prevent tissue damage.

**Figure 5.**
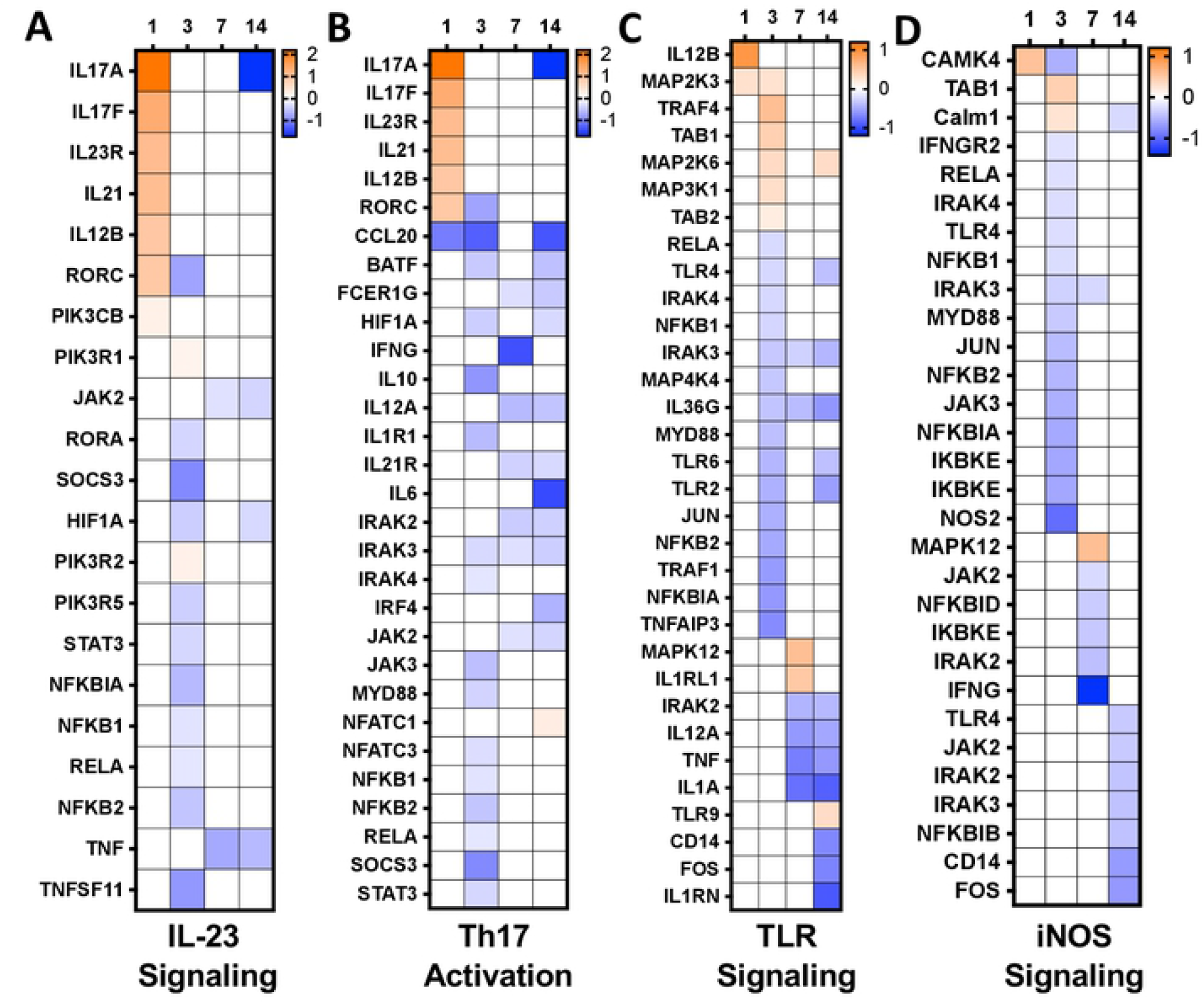
IL-27 up-regulated pro-inflammatory cytokines during the initiation of implant-associated *S. aureus* osteomyelitis and their down-regulation during chronic infection. DEGs in the **(A)** IL-23, **(B)** Th17, **(C)** TLR, and **(D)** iNOS signaling pathways are shown with log_2_ fold change on each day as heatmaps. The orange and blue colored bars indicate up-regulation or down-regulation in mice treated with rAAV-IL-27 vs. rAAV-GFP, respectively (log_2_ fold change, *p*<0.05). Pro-inflammatory cytokine coding genes such *as IL17A, IL17F, IL21,* and *IL12B* were upregulated on day-1 post-surgery. In contrast, pro-inflammatory cytokine genes such as *IL17A, IL12A, TNF,* and *IL6* were downregulated on day-14.

**Table 1.** In vitro cytokine assay with *S. aureus* infected murine bone marrow derived macrophages in the presence or absence of IL-27.

|  | PBS+*S.aureus* infection |  |  | IL-27+*S.aureus* infection |  |  |
| --- | --- | --- | --- | --- | --- | --- |
| IL-1β | 99.1 | ± | 28.7 | 90.8 | ± | 14.4 |
| IL-2 | 144.9 | ± | 2.2 | 155.2 | ± | 8.7 |
| IL-4 | 168.0 | ± | 3.3 | 174.5 | ± | 7.9 |
| IL-5 | 518.8 | ± | 9.9 | 554.4 | ± | 31.2 |
| IL-6 | 12595.4 | ± | 1263.0 | 10174.4 | ± | 2138.8 |
| IL-10 | 3721.7 | ± | 27.0 | 3915.8 | ± | 129.7* |
| IL-13 | 1354.3 | ± | 28.2 | 1397.1 | ± | 141.8 |
| IL-15 | 41387.8 | ± | 1327.2 | 42200.2 | ± | 2124.3 |
| IL-17A | 747.3 | ± | 60.4 | 778.3 | ± | 61.0 |
| IL-17F | 1079.8 | ± | 19.1 | 1099.1 | ± | 64.1 |
| IL-21 | 2279.2 | ± | 91.6 | 2439.9 | ± | 20.0* |
| IL-22 | 1827.6 | ± | 61.4 | 1859.3 | ± | 84.1 |
| IL-23 | 7492.5 | ± | 182.3 | 8042.7 | ± | 142.0 |
| IL-31 | 2454.4 | ± | 94.1 | 2672.7 | ± | 25.3** |
| IL-33 | 10965.5 | ± | 125.8 | 11595.4 | ± | 701.2 |
| IFN-γ | 164.3 | ± | 3.5 | 167.5 | ± | 8.8 |
| IFN-λ3/IL-28B | 472.9 | ± | 13.8 | 488.4 | ± | 37.1 |
| TNF-α | 500.2 | ± | 24.4 | 482.7 | ± | 9.5 |
| TNF-β | 87365.0 | ± | 1032.4 | 99471.3 | ± | 5440.6** |
| GM-CSF | 265.0 | ± | 11.1 | 281.4 | ± | 6.9 |
| MIP-3a/CCL20 | 5648.3 | ± | 247.4 | 5670.9 | ± | 37.3 |
| CD40L | 9229.2 | ± | 87.6 | 9870.0 | ± | 487.6 |

Data are presented mean ± SD (pg/mL) (N=3, \*p<0.05, \*\*p< 0.01, one-way ANOVA)

### rAAV-IL-27 treatment inhibits osteoclast formation during implant-associated osteomyelitis

μCT demonstrated that peri-implant osteolysis was decreased in infected mice subjected to rAAV-IL-27 treatment. Therefore, we hypothesized that systemic IL-27 treatment suppresses inflammatory osteoclasts to prevent bone damage during *S. aureus* osteomyelitis. IPA and gene expression analyses revealed that RANK signaling in osteoclasts was suppressed in the infected tibiae of rAAV-IL27 treated animals on days 3, 7, and 14 (**Fig. 4E, 6A**). Histopathology confirmed the suppression effect of IL-27 on osteoclasts (**Fig. 6B**), where systemic IL-27 induced significantly less osteoclast activation in trabecular bone (**Fig. 6C**).

**Figure 6.**
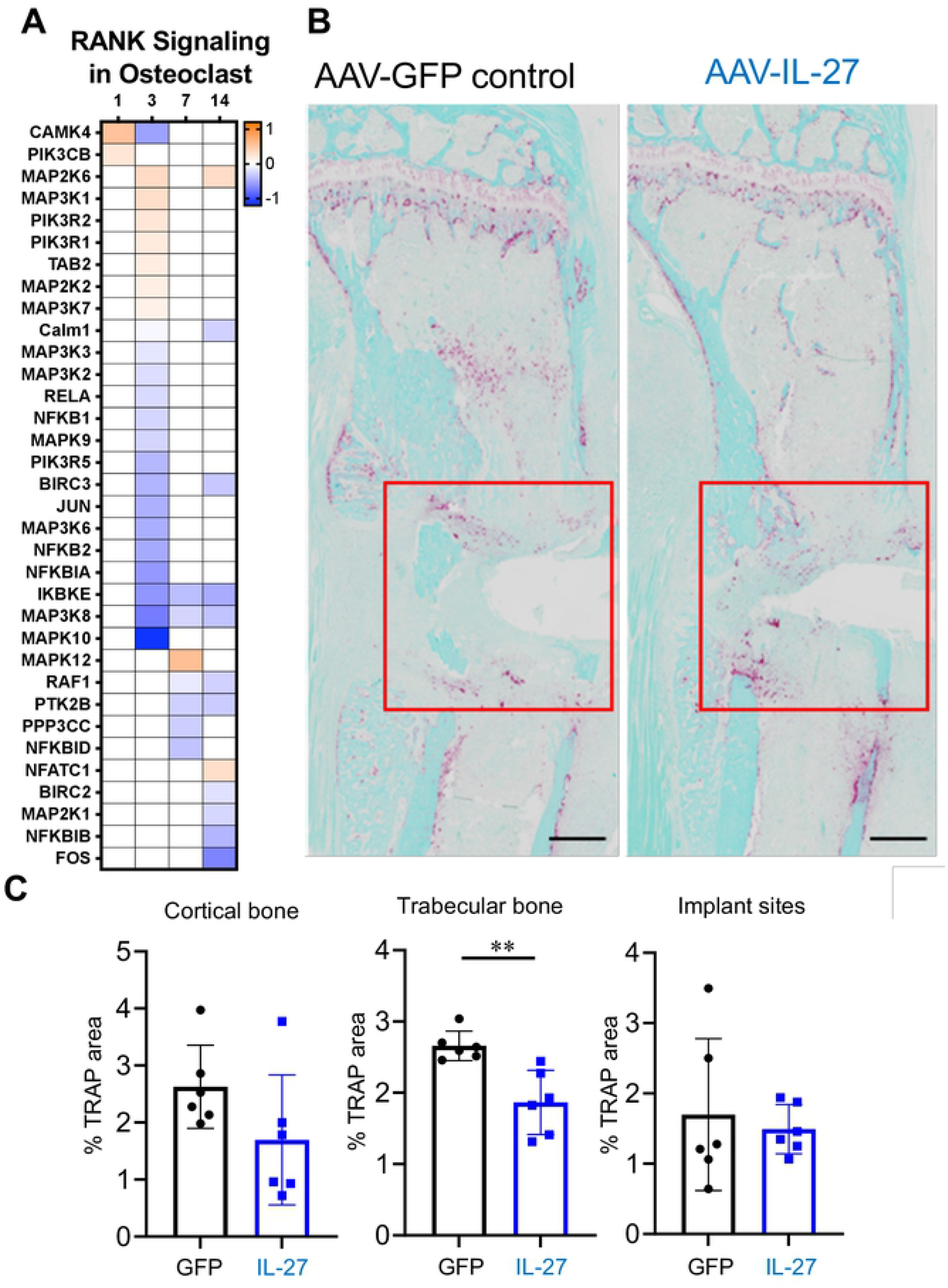
Systemic IL-27 inhibits osteoclast formation during implant-associated osteomyelitis. **(A)** Data from the IPA in Figure 4 are shown to illustrate the decrease in RANK signaling in rAAV-IL27 vs. rAAV-GFP treated infected tibiae on days 3, 7 and 14. **(B)** To confirm the gene expression data, tibiae from the mice described Figure 2 were processed for histology. Representative 2x images of tibia sections stained for TRAP (red/purple) are shown (scale bars = 500 μm). **(C)** % TRAP-stained area was quantified within the cortical bone regions, trabecular bone regions, and implant sites (red box), and the data are presented for each tibia with the mean +/− SD for the group (n=6, \*\**p*<0.05, t test).

## Discussion

Cytokines, including IL-27, are central to mounting an immune response during infection, and elucidation of IL-27 functions throughout infection is essential to our understanding of protective vs. susceptible host immunity [20]. In this study, we examined the role of IL-27 during *S. aureus* osteomyelitis as clinical studies revealed elevated serum IL-27 levels in patients with *S. aureus* bone infections. In mice, we demonstrated that IL-27/IL-27R signaling mediates bacterial clearance during the acute phase of *S. aureus* osteomyelitis, and suppresses subsequent inflammation to prevent cytokine storm and bone osteolysis during chronic infection.

A remarkable finding of our study is that serum IL-27 levels were highly diagnostic of *S. aureus* osteomyelitis in patients (AUC=0.922). Previous studies have shown that serum IL-27 levels are elevated in sepsis patients, indicating its potential as a diagnostic biomarker of sepsis [14–18, 39]. A single-center prospective study demonstrated that serum IL-27 levels could be utilized to achieve AUCs of 0.75 in patients with sepsis [16]. Though IL-27 levels immediately following septic death were 60-fold higher in patients compared to uninfected patients, we couldn’t perform AUC calculations due to the low number of septic death patients. Nonetheless, our study indicates that IL-27 could be a diagnostic marker of *S. aureus* osteomyelitis, and more extensive patient cohort studies are required to formally assess its diagnostic potential.

Systemic IL-27 delivery led to amelioration of surgical site soft tissue infection and peri-implant bone loss during the establishment of *S. aureus* osteomyelitis. However, the bacterial loads on the implant or bone were not affected by IL-27 delivery underscoring the ability of *S. aureus* to invade deep within the immune-privileged environment of bone [40]. Interestingly, reduction in abscess formation and bone osteolysis was not observed in IL-27 receptor α knock-out mice, suggesting a direct role of IL-27/IL-27R signaling on immune and bone cell functions. Similarly, Wang et al. showed that administration of recombinant IL-27 improved bacterial clearance and host survival in a rodent model of *Clostridium difficile* infection colitis [41]. Contrastingly, other studies have observed that blockade of IL-27 worsened the severity of sepsis-induced myocardial dysfunction in an endotoxic shock syndrome murine model [42]. Collectively, these studies highlight the diverse effects of IL-27 on various bacterial infections.

Transcriptome analyses of the *S. aureus* infected tibia treated with rAAV-IL-27 revealed that IL-27 is a biphasic cytokine activating pro-inflammatory pathways early during *S. aureus* osteomyelitis and suppressing them late during the chronic phase. From these observations, it is conceivable that IL-27 contributes to time-dependent changes in host immunity from acute to chronic *S. aureus* osteomyelitis. A recent study, using a murine intra-femur osteomyelitis model, demonstrated similar time-dependent changes in host response during *S. aureus* osteomyelitis using gene expression analyses [43]. We also revealed that IL-23, Th17 activation, IL-17 signaling, and pro-inflammatory IL-21 were up-regulated during early *S. aureus* infection. Collectively, these pathways contribute to the expansion of Th17 cells and induction of Th17-mediated immunity, which are crucial to host defense against bacterial infections [44, 45]. However, excessive or prolonged Th17 responses due to chronic infection cause tissue damage and autoimmune diseases [46–48]. In addition to thwarting immune responses, we observed that systemic IL-27 administration suppresses inflammatory osteoclasts to prevent bone damage during the chronic phase of *S. aureus* osteomyelitis. This is consistent with the known effects of IL-27 on inhibition of osteoclastogenesis [49–52]. Collectively, these studies add to the growing body of IL-27 literature with reported pro-inflammatory and anti-inflammatory effects on various immune cells [24, 29, 30, 53–58].

Here, we propose a schematic model of IL-27-mediated immune homeostasis during *S. aureus* osteomyelitis (**Fig. 7**). IL-27 promotes host immune reaction against *S. aureus* osteomyelitis by regulating a diverse set of immunostimulatory and immunosuppressive pathways in a time-dependent manner. At the onset of *S. aureus* osteomyelitis, IL-27 promotes the production of pro-inflammatory cytokines [29, 30, 56–58], leading to enhanced bacterial killing by macrophages and neutrophils. In contrast, at later stages of the infection, IL-27 inhibits the production of pro-inflammatory cytokines [24, 53–55] and osteoclastogenesis [49–52] to prevent cytokine storm and osteolysis. The proposed IL-27 mediated immune homeostasis model is preliminary, and warrants several further investigations. Firstly, we need to examine the immune cell repertoire in the bone marrow niche that causes IL-27/IL-27R-mediated effects during *S. aureus* osteomyelitis over time. Secondly, we need to understand IL-27’s role in preventing cytokine storm and internal organ tissue damage during chronic *S. aureus* osteomyelitis in a more relevant osteomyelitis sepsis murine model. Humanized mice, which are more susceptible to MRSA osteomyelitis-induced sepsis, may be better suited for these studies [59]. Finally, we need to assess how systemic IL-27 inhibits bone osteolysis by suppressing RANKL-mediated osteoclastogenesis. These studies will further our understanding of IL-27/IL-27R signaling during *S. aureus* osteomyelitis.

**Figure 7.**
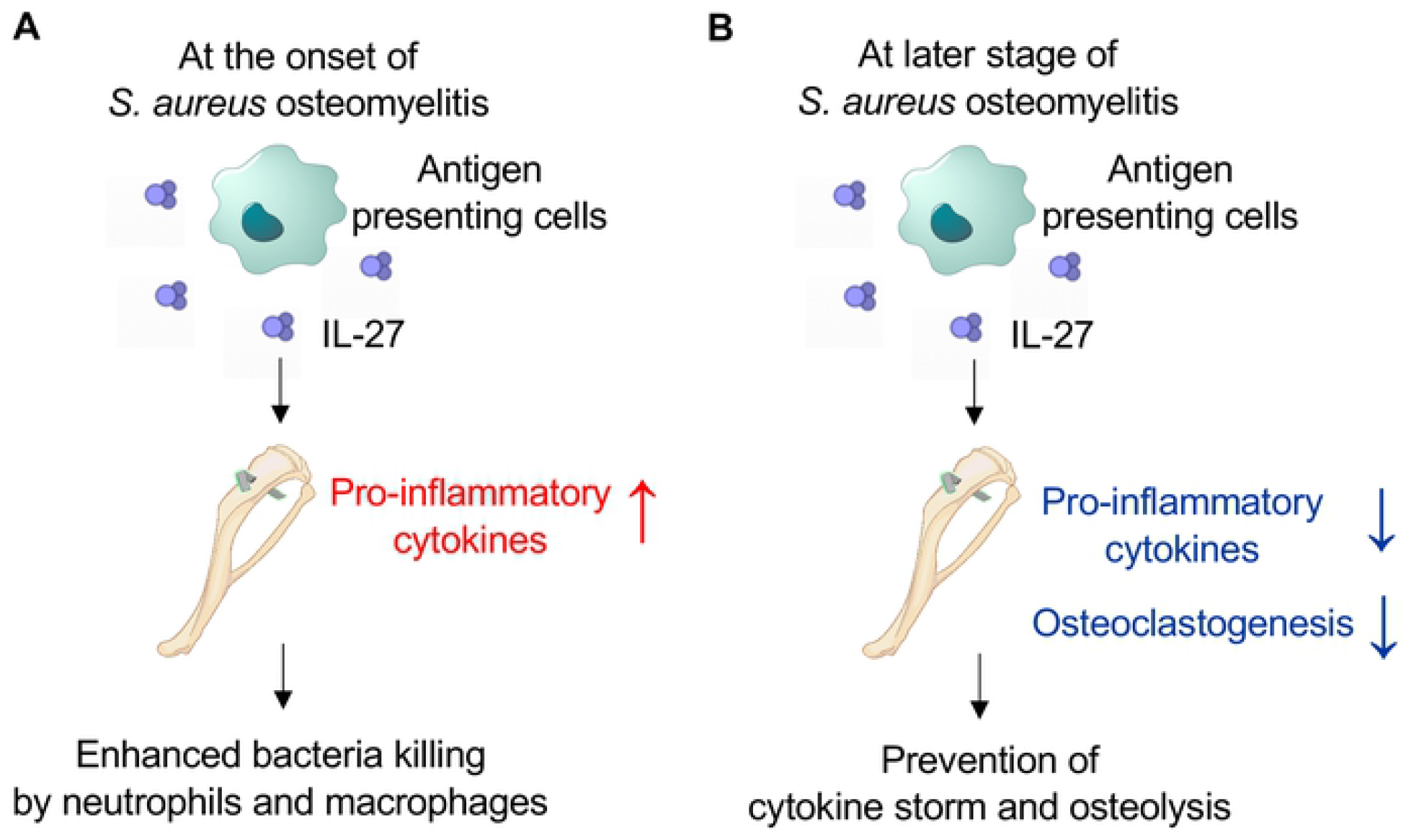
Schematic model of IL-27 mediated immune homeostasis during *S. aureus* osteomyelitis. Here, we propose a model of IL-27-mediated immune homeostasis during *S. aureus* osteomyelitis in which IL-27 promotes host immune reaction against *S. aureus* osteomyelitis by regulating its reported diverse immune-activation and immune-suppression effects in a time dependent manner. **(A)** At the onset of *S. aureus* osteomyelitis, IL-27 promotes production of pro-inflammatory cytokines, leading to enhanced bacteria killing by macrophages and neutrophils. **(B)** In contrast, at later stages of *S. aureus* osteomyelitis following acute reaction, IL-27 decreases production of pro-inflammatory cytokines and osteoclastogenesis, which prevents cytokine storm and osteolysis.

## Materials and Methods

### Bacterial strains

Methicillin-resistant *S. aureus* (USA300 LAC) was used for all in vitro experiments, and a bioluminescent strain of USA300 (USA300 LAC::lux) was used for all in vivo experiments as previously described [32–34, 36, 59].

### Ethics Statement and Patient Enrollment

Serum samples were collected from *S. aureus* osteomyelitis patients (n=23) and uninfected patients undergoing elective total joint replacement (n=10). Additionally, serum samples were collected immediately post-mortem in patients that succumbed to *S. aureus* osteomyelitis sepsis (n=5). All recruited patients were either part of an international biospecimen registry (AO Trauma Clinical Priority Program (CPP) Bone Infection Registry) [60] or clinical studies conducted at the Virginia Commonwealth University. Patients were recruited with local IRB approvals at various institutions, and patient information was collected in a REDCap database managed by AO Trauma and VCU data management administrators. Laboratory investigators had access only to de-identified clinical data, which was provided on request by the data management teams. All ex vivo and in vivo mouse infection studies were performed at the University of Rochester in accordance with protocols approved by the Institutional Animal Care and Use Committee at the University.

### Luminex-based cytokine measurements

Serum IL-27 levels were determined in patients via Luminex assay using the Milliplex xMAP Multiplex Assay (MilliporeSigma) according to the manufacturer’s instructions. Primary bone marrow-derived murine macrophages (BMDMs) were pretreated with PBS or murine IL-27 (50 ng/ml from BioLegend) for 24 hours and then infected with *S. aureus* at MOI = 10 in the presence or absence of murine IL-27 (50 ng/ml) for 24 hours. Subsequently, the cell culture supernatants were harvested from these cells to measure the following cytokines using a multiarray Milliplex xMAP murine cytokine Magnetic Bead Panel according to manufacturer’s instructions: CD40L, GM-CSF, IFN-γ, IL-1β, IL-2, IL-4, IL-5, IL-6, IL-10, IL-13, IL-15, IL-17A, IL-17F, IL-21, IL-22, IL-23, IFN-λ3/IL-28B, IL-31, IL-33, MIP-3α/CCL20, TNF-α, and TNF-β.

### In vitro IL-27 induction assay in macrophages

Primary BMDMs from 12-week-old C57BL/6 mice (Jackson Laboratory) were isolated from femur and tibia. After dissection of the femur and tibia from mice, bones were washed in RPMI + 10% FBS, 1% HEPES, and 1% anti-microbial/anti-mycotic (R10) media before disinfection using 70% ethanol. After disinfection, the long bones were cut on both ends, and marrow was flushed using a 23G needle and resuspended in R10 media. After spinning cells down at 500g for ten minutes, the isolated cells were resuspended in a red lysis buffer to remove red blood cells. Cells were then resuspended again in R10 media with mouse colony-stimulating factor (M-CSF) (25ng/mL) and plated at 5 × 10^6^ cells/plate for 6 days. Subsequently, BMDMs were differentiated with PBS, murine IFN-γ (50 ng/ml from PeproTech) or murine IL-4 (20 ng/ml from PeproTech) in R10 for 24 hours to generate M0, M1, and M2 cells respectively [61]. These cells were then infected with *S. aureus* USA300 at MOI = 10 for 24 hours, and subsequently, the cell culture supernatant was harvested to examine IL-27 secretion using the Mouse IL-27 Uncoated ELISA kit (Invitrogen).

### Reactive nitrogen species induction in murine macrophages

Murine BMDMs were pretreated with PBS or murine IL-27 (50 ng/ml from Biolegend) for 24 hours, and then stimulated with or without LPS (100ng/ml from MilliporeSigma) to induce reactive nitrogen species production [62], which is important for host defense against bacterial infection [63]. Stimulation experiments were performed on BMDMs with or without murine IL-27 (50 ng/ml) for 24 hours after pretreatment. Subsequently, nitrite concentrations in cell culture supernatant were determined via Griess reaction assay kit (R&D Systems).

### Transwell Chemotaxis assay

HL-60 cells (ATCC) were differentiated into granulocytes using 100mM dimethylformamide (DMF) (MilliporeSigma), placed on top of Boyden chambers, and chemotaxis assay was performed according to manufacturer’s protocol (MilliporeSigma QCM™ Chemotaxis 5 μm 24-Well Cell Migration Assay kit). Briefly, 1×10^6^ cells/chamber were subjected to chemotaxis in RPMI media with or without the chemoattractant (human IL-27 (500 ng/ml from PeproTech) or N-formyl-methionyl-leucyl-phenylalanine (fMLP) (800 ng/mL from MilliporeSigma) as positive control)) placed below the chamber, and incubated for 1 hour at 37 °C. Post incubation, cell migration from the chambers was enumerated as relative fluorescence units (RFUs) according to the manufacturer’s instructions.

### RecombinantIL-27-expressing adeno-associated virus vector (rAAV-IL-27) administration

To get sustained exogenous IL-27 expression, mice were subjected to intramuscular administration of recombinant murine IL-27-expressing AAV (0.5 × 10^12^ genome copies/mouse, Vector Biolabs) 7 days prior to *S. aureus* septic surgery [64]. Mice intramuscularly infected with AAV expressing recombinant GFP (0.5 × 10^12^ genome copies/ mouse, Vector Biolabs) were used as controls.

### Implant-associated MRSA osteomyelitis in mice

C57BL/6 and IL-27Rα deficient mice (IL-27Rα^−/−^) in the C57BL/6 background used in the study were purchased from Jackson Laboratories and maintained at the University of Rochester animal facilities. All in vivo *S. aureus* challenge experiments in mice utilized our well-validated transtibial implant-associated osteomyelitis model [32–34, 36, 59]. Briefly, L-shaped stainless-steel implant was contaminated with USA300 LAC (5.0 × 10^5^ CFU/mL) grown overnight, and surgically implanted into the tibia of 8-week-old female C57BL/6 mice from the medial to the lateral side. Longitudinal body weight change and bioluminescent intensity at infection site were evaluated, and terminal assessment of CFU (implant, surgical site soft tissue and tibia), peri-implant osteolysis (high-resolution μCT imaging), biofilm formation on the implant (Zeiss Auriga SEM imaging), and histopathology were performed on day 14 post-septic surgery as described previously [32–34, 36, 59]. Murine infection studies were performed three independent times, and the resulting data was pooled from these experiments.

### RNA sequencing of MRSA infected tibia

C57BL/6 were intramuscularly injected with rAAV-IL-27 or rAAV-GFP and then challenged with an *S. aureus* contaminated transtibial implant as described above. Infected tibiae were collected on days 1, 3, 7, and 14 post-surgery for RNA sequencing. Tibiae were pulverized in liquid nitrogen (−196°C) and homogenized using Bullet Blender Gold (Next Advance). Collection of Total RNA from homogenized tibia was performed using TRIzol extraction (ThermoFisherScientific) and RNeasy Mini Kits (Qiagen). Contaminating genomic DNA was removed using TURBO DNase (ThermoFisherScientific). The TruSeq Stranded Total RNA Library Prep Gold (Illumina) was utilized for next-generation sequencing library preparation per the manufacturer’s instructions. The libraries were sequenced with the Illumina NovaSeq6000 platform (Illumina). Quality filtering and adapter removal were performed by fastp version 0.20.0 [65] using the following parameters: “--in1../$(SAMPLE)_R1.fastq.gz --out1 clt_$(SAMPLE)_R1.fastq.gz --length_required 35 --cut_front_window_size 1 --cut_front_mean_quality 13 --cut_front --cut_tail_window_size 1 --cut_tail_mean_quality 13 --cut_tail -w 8 -y -r -j $(SAMPLE)_fastp.json”. The remaining high quality processed reads were then mapped to the *Mus musculus* genome reference (GRCm38.p6) with STAR version 2.7.0f [66] using the following parameters: “--twopassMode Basic --runMode alignReads --genomeDir $(GENOME) --readFilesIn $(SAMPLE) --outSAMtype BAM Unsorted --outSAMstrandField intronMotif --outFilterIntronMotifs RemoveNoncanonical”. The mapped reads were counted within the GRCm38.p6 gene annotations using the featureCounts read quantification program in Subread version 1.6.4 [67]. Then, the differential expression analyses and data normalization were performed using DESeq2 version 1.22.1 [68] within the R version 3.5.1 with a p-value threshold of 0.05 on each set of raw expression measure. Subsequent bioinformatics analyses including Canonical Pathway Analysis and Regulator Effect Network Analysis were performed using Ingenuity Pathway Analysis (IPA; Qiagen) for each time-point. All generated sequence data have been submitted to Gene Expression Omnibus with accession number GSE168896.

### Statistics

For statistical analyses, the non-parametric Kruskal-Wallis test, one-way ANOVA, repeated measures two-way ANOVA, and unpaired student’s t-test were utilized to assess significance between experimental groups. Data were presented as mean ± standard deviation. A *p* value less than 0.05 was considered significant.

## Acknowledgments

The authors would also like to thank Drs. Chad Galloway and Elysia A. Masters for their technical assistance. The authors would like to thank the Electron Microscope Shared Resource Laboratory, Genomics Research Center, and the Histology, Biochemistry, and Molecular Imaging Core in the Center for Musculoskeletal Research at the University of Rochester Medical Center.

## Funding

This work was supported by NIH NIAMS P30 AR069655 pilot grant (GM) with additional support from NIH NIAMS P50 AR072000 (EMS), P30 AR069655 (EMS), and AO Trauma Clinical Priority Program (EMS, SLK).

## Conflict of interest statement

The authors have declared that no conflict of interest exists.

## Data availability

All RNA sequence data have been submitted to the Gene Expression Omnibus with accession number GSE168896.

---

**Supplemental Figure 1.**
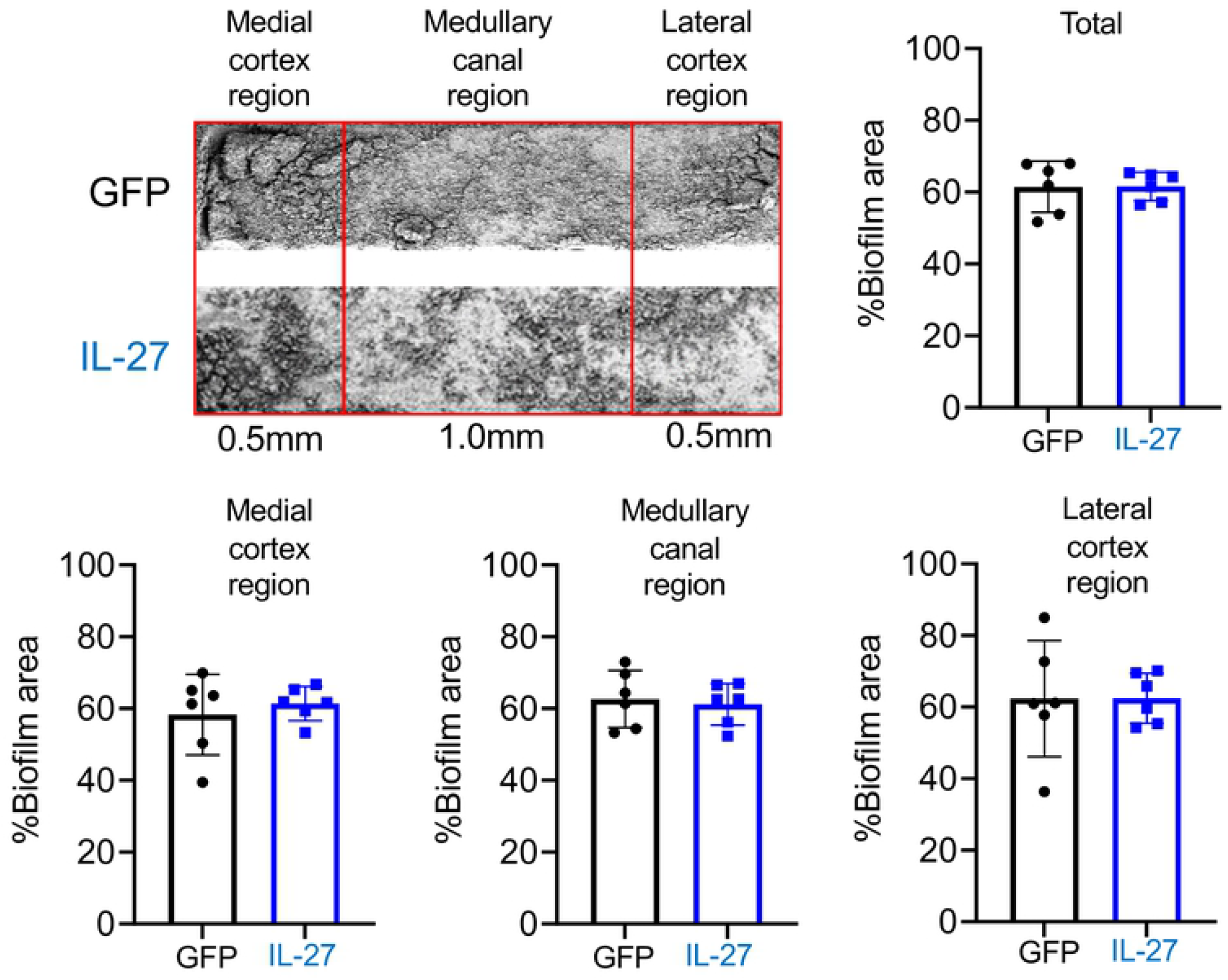
Systemic IL-27 does not affect biofilm formation on the implant during *S. aureus* implant-associated osteomyelitis in vivo. Mice were intramuscularly injected with rAAV-IL-27 or rAAV-GFP and then challenged with a MRSA (USA300 LAC::lux) contaminated transtibial implant as described in Figure 2. Biofilm formation on the implant was determined via SEM processing and imaging after euthanasia on day 14 post-op. No difference was detected in %biofilm formation area on implant between rAAV-IL-27 and rAAV-GFP challenged mice (n=6).

**Supplemental Figure 2.**
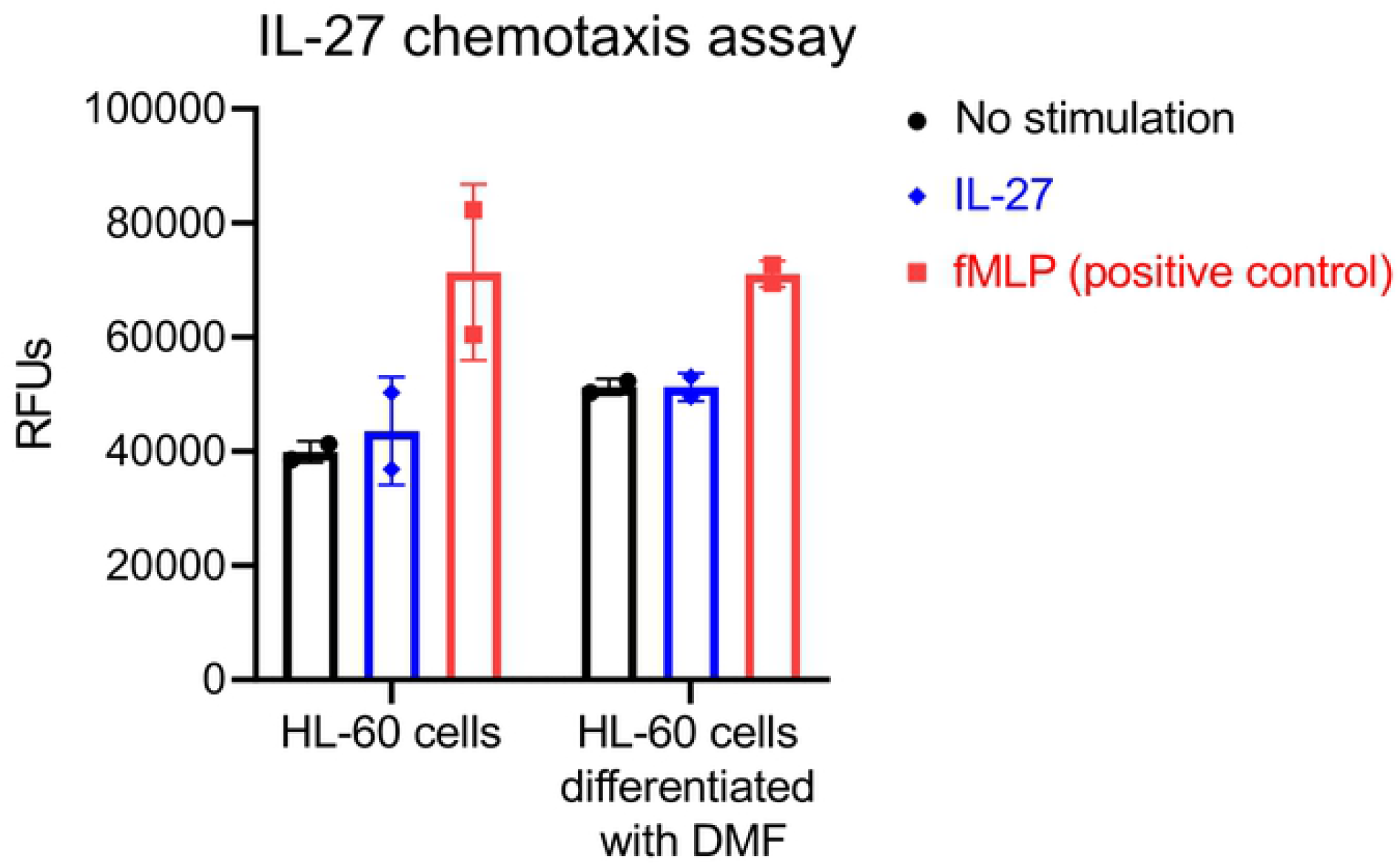
IL-27 does not stimulate myeloid cell chemotaxis. HL-60 cells were differentiated 7 days in the presence or absence of dimethylformamide (DMF) (9 μg/ml), and then placed in Boyden chambers. Cell culture media with or without IL-27 or fMLP (positive-control) was placed in the well below the chamber and incubated for 1 hour. Subsequently, cells which migrated in each well were stained with fluorescent dye and signal intensity was evaluated using a fluorescent plate reader (n=2). No difference in chemotactic activity of granulocytes was observed between the experimental groups.

**Supplemental Figure 3.**
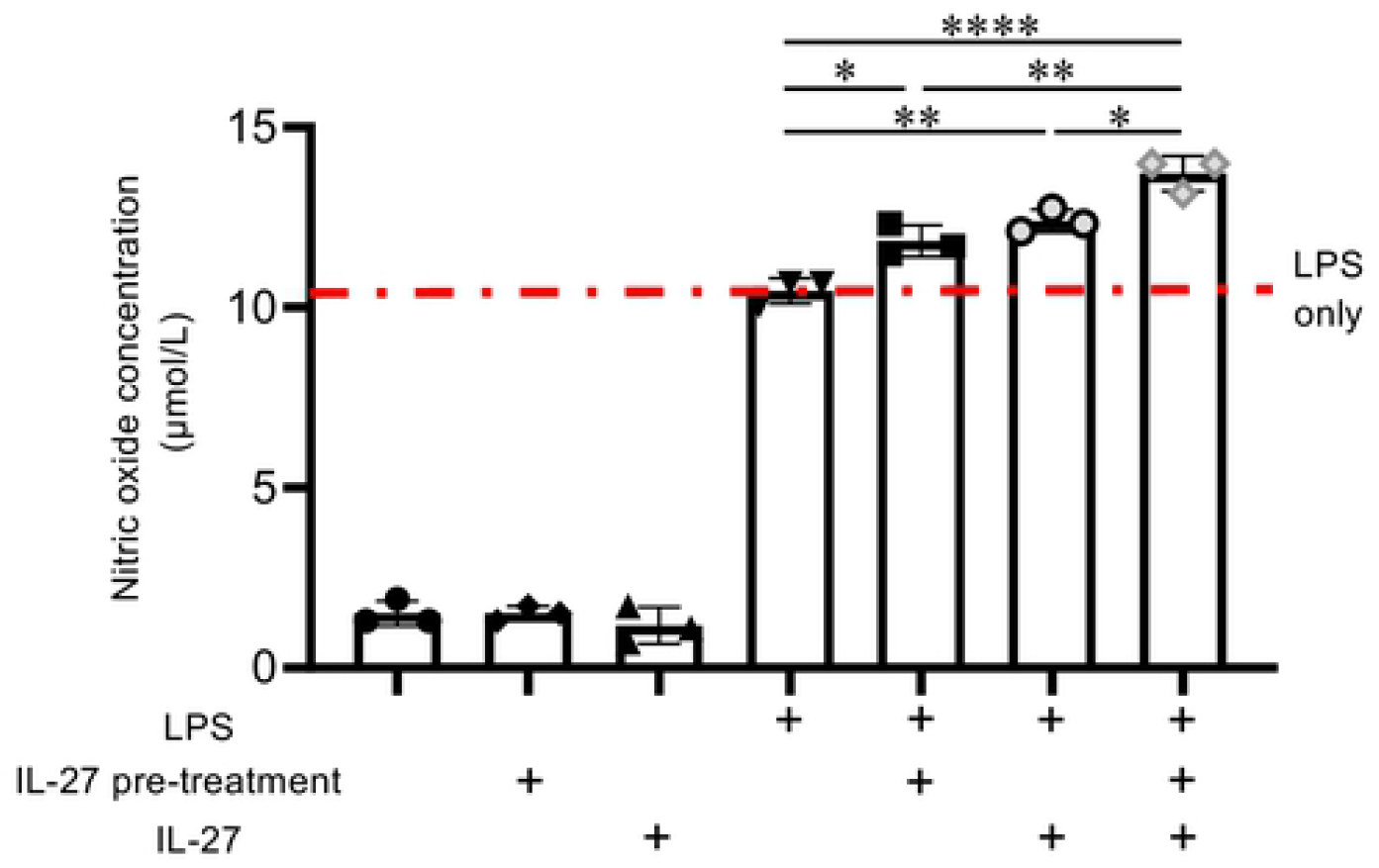
IL-27 enhances LPS-induced NO^−^ production by macrophage cultures. Primary bone marrow derived murine macrophages were pretreated with PBS or IL-27 (50 ng/ml) for 24 hours, and then stimulated with LPS (100 ng/ml) in the presence or absence of IL-27 (50 ng/ml) for 24 hours. Nitrite levels in the culture supernatant was determined via Griess reaction assay, and the data from each experiment are presented with the mean +/− SD for the group (n=3, \**p*<0.05, \*\**p*<0.01, \*\*\*\**p*<0.0001 by one-way ANOVA).

**Supplemental Table 1.** Top regulatory effect networks of DEGs in mice treated with rAAV-IL-27 versus rAAV-GFP.

|  |  |  |  |  |  |
| --- | --- | --- | --- | --- | --- |
|  |  | Consistency score | Regulators | Target molecules in dataset | Functions |
| Day 1 | 1 | 14.432 | HSP90B1, HBB, EGR2, BCL2, TNFRSF4, LTA | BCL2A1, CXCL13, CCL20, CXCL9, CXCL6, IL7R, AIF1, IL21, IL17A, IL12B, VCAM1, IL17F, RORC, GZMB | Inflammation of body cavity, Inflammation of organ, Inflammation of absolute anatomical region, Inflammation of respiratory system component: **Predicted activation** |
|  | 2 | 8.05 | NR1H | IL12B, PTGES, MERTK, ABCG1, CD5L | Inflammation of absolute anatomical region, Inflammation of body cavity, Inflammation of respiratory system component, Cell death of immune cells, Apoptosis of blood cells: **Predicted activation** |
| Day 3 | 1 | 11.333 | MYD88, TICAM1, GFI1, IL33, IGE,NFKB, NFKB1, NOD2, FOS, IKBKB | IL1R1, EDNRB, HDC, IL10, ICAM1, MDM2, SMAD3, C5, GRN | Increased levels of creatinine: **Predicted suppression** |
|  | 2 | 10.436 | JUNB, SMARCA4, STAT3, TNFSF11, IL1, IL1A, OSM, CHUK, IL6R, CD40LG, TGFB2, RELA | IFI16, CSF3, NOS3, IL11, IGF2, TNFRSF11B, RB1, SMAD3, NOTCH1, HES1 | Increased activation of alkaline phosphatase: **Predicted suppression** |
| Day 7 | 1 | 3.175 | TICAM1 | IL12A, IRF7, CCL5, IFNG, TNF, IL1A, DDX58, CH25H, FPR1, ICAM1, CCRL2, IKBKE | Cell movement of blood cells: **Predicted suppression** |
|  | 2 | 3.175 | TICAM1 | IL12A, IRF7, CCL5, IFNG, TNF, IL1A, DDX58, CH25H, FPR1, ICAM1, CCRL2, IKBKE | Leukocyte migration: **Predicted suppression** |
| Day 14 | 1 | 4.025 | IL1B | CCR7, ITGAM, OSM, ICOSLG/ LOC102723996, CCL20, LIF, IL6, SELE, CD44, LCP1, TLR4, CXCR4, TNF, IL1A, CD80, IL17A, FAS, CCL4, MIF, ICAM1 | Adhesion of mononuclear leukocytes: **Predicted suppression** |
|  | 2 | 3.873 | IL17A | TGFA, HBEGF, LIF, COL1A1, SLC2A1, IL6, FOS, TLR4, AREG, TNF, IL1A, TIMP1, CSF3, FAS, IL36G | Proliferation of connective tissue cells: **Predicted suppression** |

## Notes

### Competing Interest Statement

The authors have declared no competing interest.

